# The Sikkim mouse (*Mus pahari*) exhibits distinct spatial, circadian, and social behaviors compared to laboratory mice

**DOI:** 10.1101/2025.11.13.688123

**Authors:** Caleb C. Vogt, Michael J. Sheehan

## Abstract

While the laboratory mouse is one of the most studied organisms on the planet, comparative research on the spatial and social structures of closely related *Mus* species remains limited. Here, we characterize the spatial, circadian, and social behavior of the Sikkim mouse (*Mus pahari*) wild-derived inbred strain PAH/Eij in comparison to the laboratory mouse strain C57BL/6J (*Mus musculus domesticus*, shortened to ‘B6’). Using a common garden approach, we monitored mice in replicate mixed-sex group social behavior trials within an indoor mesocosm using radiofrequency identification (RFID) antennae. *M. pahari* exhibited markedly reduced spatial exploration and highly stereotyped circadian activity that was strongly coupled to the dark phase of the light-dark cycle compared to B6 mice. Most strikingly, *M. pahari* displayed distinctive social behaviors characterized by strong male-male gregariousness and enhanced overall social tolerance, contrasting sharply with the relatively less social nature of B6 mice. These results demonstrate the genetic influence on social organization within the *Mus* genus and identify unique socio-spatial phenotypes in *M. pahari*, highlighting the considerable potential of this strain and species as a novel model for social behavior research.

## INTRODUCTION

Social behavior is a fundamental aspect of species’ biology, and understanding the factors shaping its expression and evolution remains a central goal in behavioral ecology^1,2^. Group living and social organization profoundly impact individual fitness by influencing resource acquisition, reproductive access, and survival^3,4^. Comparative studies of closely related organisms can provide valuable insights into the neural, endocrine, genetic, and evolutionary forces shaping the expression of specific social behavior phenotypes^5–8^. The comparative approach is particularly powerful when applied to clades containing well-established model species, such as the genus *Mus*, which includes 38 species across four subgenera, including the well-studied lab mouse^9,10^.

In recent years, researchers have increasingly turned to diverse rodent species like blind mole rats^11^, prairie voles^12^, African striped mice^13^, spiny mice^14^, and deer mice^15^ to study the behavioral, genetic, and neurobiological mechanisms underpinning complex social behaviors (**Fig. 1A**). However, relatively few studies have explored the potential of closely related species within the *Mus* genus, which may offer advantages both in terms of expressing novel phenotypes and leveraging the extensive genetic and genomic resources developed for the closely-related house mice. For instance, *Mus spicilegus*, the mound-building mouse, exhibits strong mating pair-bonds, significant paternal care, and cooperative behaviors in building large mounds of vegetation for overwintering^16–18^. *Mus spretus*, the Algerian mouse, has been used in comparative studies of aggression and territorial behavior^19^, and, along with *Mus caroli*, has contributed to our understanding of genome evolution and speciation in the *Mus* genus^20,21^.Our study adds to this growing body of comparative work within the *Mus* genus by examining *Mus pahari*, a species that has received relatively little attention in behavioral research despite its close evolutionary relationship to the model lab mouse, the availability of a fully sequenced genome, and a wild-derived inbred strain. *M. pahari*, also known as the Sikkim mouse or Gairdner’s shrewmouse, is found in northeastern India, Thailand, Bhutan, Myanmar, Vietnam, and southern China^10^. *M. pahari* diverged from the *M. m. domesticus* lineage between 5-7.6 million years ago^9,22,23^ (**Fig. 1A**), representing one of the more evolutionarily distant *Mus* species for which laboratory strains are available. Species in this subgenus typically appear shrew-like with long noses, short ears, small eyes, and bluish gray velvet/spiny coats^24,25^ (**Figs. 1B & S1B**), morphological traits that hint at a potentially distinct ecological niche relative to the house mouse. While descriptions of their biogeography and behavior are scant, Marshall (1977) reported that the species can be found in tall grassy clearings within mountain rain forests above 1800 meters^24^. Captive wild-caught *M. pahari* are described as highly active (e.g. constantly exercising on a running wheel even during the daytime hours), and capable of emitting “high piping calls, uttered rapidly and at descending pitch”^24^. Details of their population dynamics are likewise scarce, though one report found a female-biased sex ratio in one wild sample^26^.

**Figure 1.**
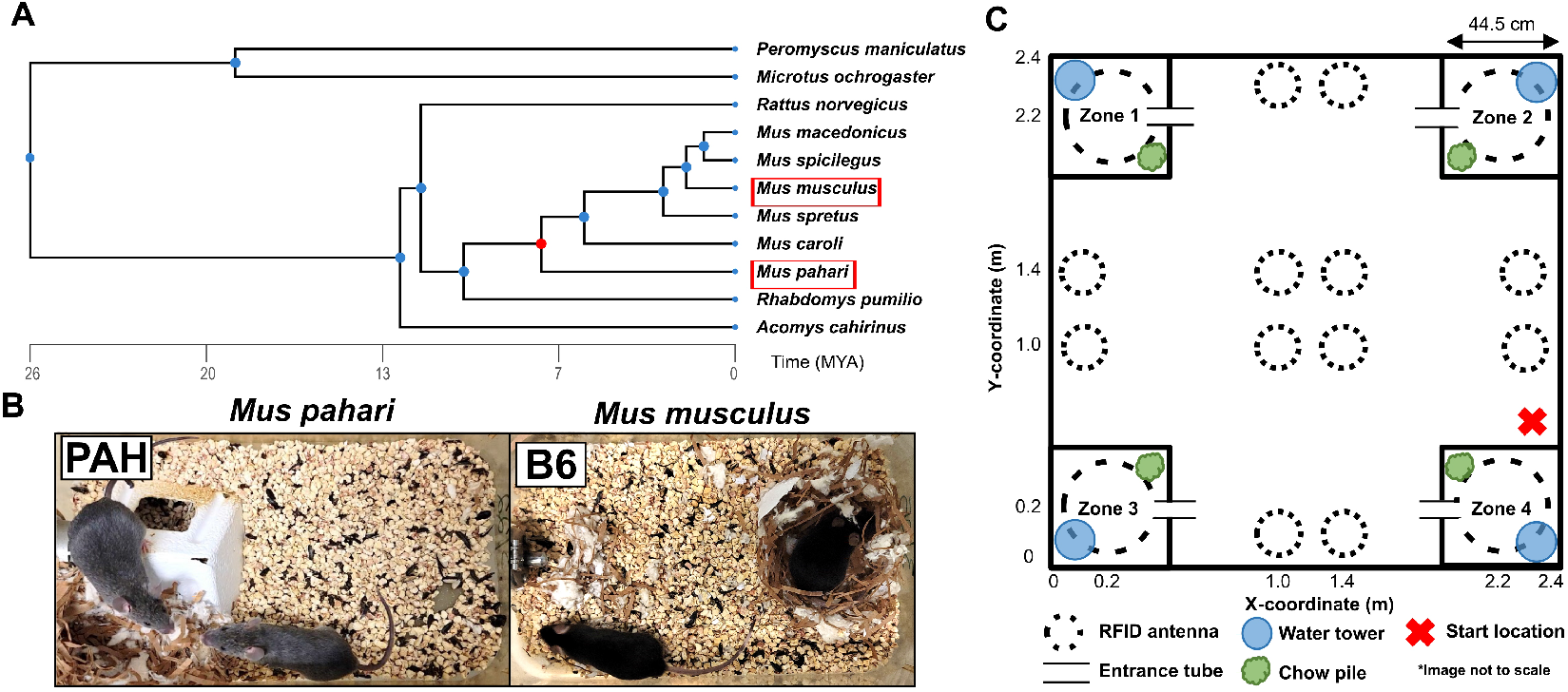
Comparison of Mus pahari and Mus musculus domesticus in a common garden mesocosm. **A**. Phylogenetic tree showing evolutionary divergence of *M. pahari, M. m. domesticus* (C57BL/6J), and other rodent species used in laboratory social behavior research. **B**. Representative images show the distinctive morphology of *M. pahari* (PAH/EiJ) with its shrew-like appearance compared to the familiar *M. m. domesticus* (C57BL/6J). Tree generated using TimeTree software. **C**. Schematic of the indoor mesocosm arena (2.4 × 2.4 m) used for the common garden behavioral trials. RFID antennas (n = 16, black dotted circles) were positioned throughout the arena to monitor spatial movements and social behaviors of the mice. Resource zones (n = 4, black squares in corners) provided ad libitum food and water. Each trial included 4 males and 4 females of the same genotype. Three independent trials were conducted for each genotype (n = 24 mice/genotype, 48 mice total) over five days each to compare patterns of spatial, circadian, and social behaviors.

The *M. pahari* genome provides a valuable resource for comparative genomic studies within the *Mus* genus and for investigating the genetic basis of novel phenotypes divergent from traditional laboratory strains^21,27–30^. For example, the inbred laboratory strain of *M. pahari* used in this study (PAH/EiJ, The Jackson Laboratory) displays skin-fragility, particularly affecting the tail, a feature not present in other common mouse strains, although we note that this attribute was not reported in earlier descriptions of wild-caught samples of *M. pahari*^24,25,31,32^. Thus, it remains a possibility that this trait may be a result of inbreeding and may not be representative of wild *M. pahari* populations.

Here we characterize key features of spatial behavior, circadian activity, and social organization in mixed-sex groups of inbred *M. pahari* and compare them to the inbred C57BL/6J lab strain of house mice. By examining these closely related species under modestly complex but controlled social and physical conditions, we aim to provide a direct comparison of behavioral differences between species. While we acknowledge that the use of an inbred *M. pahari* strain may not capture the full behavioral repertoire of wild populations, this approach allows us to control for within-strain genetic variation and focus on species-level differences. Additionally, while prior research highlights many behavioral and physiological differences between lab mice and their wild house mouse relatives^33–37^, work on social behavior organization specifically has found many similarities between lab and wild mice as well^38–40^. Our findings reveal that *M. pahari* exhibits reduced movement, strict circadian activity, and highly gregarious social behavior compared to C57BL/6J. These differences suggest the potential of *M. pahari* as a novel model system for studying the genetic and neurobiological mechanisms underlying complex sociality within the well-characterized *Mus* genus, while also highlighting the need for future studies examining the behavioral ecology of wild *M. pahari* populations.

## METHODS

### Mice

We selected adult (70-140 days old), reproductively inexperienced male and female mice from laboratory colonies of *M. pahari* and *M. m. domesticus* C57BL/6J. *M. pahari* (PAHARI/EiJ; #002655) and C57BL/6J (#000664) mice were obtained from The Jackson Laboratory (Bar Harbor, Maine, USA) and subsequently bred and maintained in our colony at Cornell University. While we note that the PAH/EiJ strain is no longer commercially available from the Jackson Laboratory at the time of this writing, M.J.S. maintains a breeding colony and can be contacted regarding availability of these mice. All animals in this study were bred and maintained under the same climate-controlled conditions in an Animal Care facility with a 14:10 light:dark cycle and were provided food and water *ad libitum*. Colony cages were identical across genotypes, containing corn cob bedding, cardboard huts, cotton nestlets, and running wheels for enrichment and exercise. Mice were minimally handled and were moved using transfer cups for all procedures to reduce handling stress. All experimental procedures were approved by the Cornell University Institutional Animal Care and Use Committee (IACUC: Protocol #2015-0060) and adhered to guidelines established by the U.S. National Institutes of Health and the Association for the Study of Animal Behavior^41^.

### Mesocosm enclosure and behavioral trials

We constructed a semi-natural mesocosm arena using ½”-thick polyvinyl chloride (PVC) boards, creating a 2.4 × 2.4-meter (4.84 m^2^) enclosure with walls approximately 0.75 m tall to prevent escape. For comparison, this arena is more than three times larger than the semi-natural mesocosms used in other recent studies of house mouse social organization and behavior^40,42^. The arena was elevated 0.3 m on a painted, pressure-treated lumber frame, allowing access to the underside of the arena for RFID antenna installation. The arena was arranged in four quadrants that contained identical enrichment materials positioned in the same configuration (**Figs. 1C, S1A**). Each corner of the arena housed a resource zone – a PVC box (44.5 × 44.5 × 50 cm) accessed through a single 10 cm PVC tube entrance. These zones contained a running wheel for exercise, plexiglass hut for shelter, and a water tower (~1 liter) and a pile of loose lab chow providing mice with food and water *ad libitum*. Red acrylic sheets covered the resource zones, providing shade during the light cycle. We spread wood chip bedding approximately 1 cm deep across the entire arena floor, allowing mice to create nests and providing absorbent substrate.

A total of 16 RFID antennae were mounted to the underside of the floor of the arena using single-sided Velcro tape in a grid formation with precise coordinates measured from the center point of the antenna. Twelve 6” diameter RFID antennae were mounted in the center of the arena, while an additional four 10” diameter RFID antennae were used to monitor the four resource zones (one antenna per zone). RFID antennae were connected to a centralized data acquisition unit (BioMark, Small Scale System, Boise, ID, USA). Each antenna had a read range of approximately 5-10 cm above the plane of the antenna, which allowed for the detection of mice moving above the antenna through the PVC floor of the arena. The antenna system scanned at approximately 2-3 Hz continuously for the duration of each trial. Each resource zone was monitored via a single infrared camera mounted in the acrylic cover of the zone, while a single fish-eye lens infrared camera mounted above the mesocosm arena monitored the entirety of the enclosure space.

Trials occurred between October 2021 and April 2022. Four male and four female virgin adult mice were used in each trial replicate (n = 3 trials/genotype). Each trial consisted of a five-day uninterrupted experience within the mesocosm enclosure. Prior to the start of the experiment, mice were same-sex sibling housed with up to three other cage mates. Within a trial, no males were siblings, but up to two females could be siblings. At least six days prior to the start of a trial, mice were briefly anesthetized with isoflurane (3-5%) and subcutaneously implanted with dual passive integrated transponder (PIT) tags (BioMark, Boise, ID, USA) in the dorsal flank and periscapular region. After implantation, male mice were individually housed in new cages while females were returned to their previous cage with cage mates. Males and females were subsequently transferred to a new temperature and humidity-controlled colony room which was on a 14:10 shifted light:dark cycle (dark period: 1200-2200 hours), matching the conditions in the mesocosm procedure room next door.

One hour prior to the start of the trial, we weighed each mouse and confirmed their identities by scanning the implanted PIT tags with a handheld RFID reader (BioMark, HPR Lite). Trials were initiated immediately following lights off at 1200h on the first day by individually removing mice from their cages using plastic transfer cups which were gently placed on the floor the enclosure, allowing the mice to freely walk out into the arena. Mice were sequentially released in a random order over the course of approximately five minutes. The trials were then left undisturbed for five days. At the end of the fifth day between 1000-1400 hours, mice were gently encouraged to enter the resource zone spaces after which the PVC entrance tube was blocked to prevent escape. Mice were then placed in transfer cups, weighed, and moved to another room where they were euthanized.

Between trials, we thoroughly cleaned the arena by removing all bedding and sanitizing all surfaces with 70% ethanol, with a second 70% ethanol scrub the following day. We washed all enrichment materials using high-pressure, high temperature cage washing equipment. The arena was prepared with fresh materials at least two days before the next trial.

### Circadian rhythm

Circadian activity was quantified using identified individual transition events between RFID antennas as a proxy of gross locomotor activity. Because antennas were non-adjacent within the arena (**Fig. 1C**), a transition event between two distinct antennas necessarily reflected physical movement by the mice across space. Thus, antenna transitions served as an index of overall locomotor activity and wakefulness throughout the light-dark cycle.

For each individual, we calculated the total number of antenna transitions to generate continuous activity time series and visualized this activity as actograms spanning successive 24-h periods. The proportion of total daily transitions occurring during the dark period was used as a measure of nocturnality. To further assess rhythmicity, we performed Lomb-Scargle periodogram analyses on the normalized transition time series for each animal. This method identifies dominant periodicities in irregularly sampled or incomplete datasets, making it well suited for RFID-based activity records.

### Social behavior

Social behavior was quantified by estimating the time mice spent in spatiotemporal association at RFID antennae following previously established methods^38,43^. Because RFID detections represent discrete spatial locations within the arena, simultaneous detections of multiple mice at the same antenna indicated physical co-presence and were used to infer social associations.

Raw RFID reads were first grouped into antenna visit bouts for each mouse by combining sequential detections separated by ≤ 300 s. Spatiotemporal association bouts were then defined as overlapping visit bouts of two or more mice at the same antenna, thereby capturing the onset and duration of social grouping events. The cumulation duration of all such associations per individual to estimate total social time, and bouts were further parsed by group composition (male-male, female-female, and male-female).

RFID-detected associations could occur either within resource zones or in open areas of the enclosure. The majority of social grouping occurred within the resource zones, but all antenna locations were included in analyses. Sociality was summarized for each individual as the percentage of total observed time spent with ≥1 conspecific.

### Data acquisition and statistical analyses

We offloaded RFID data daily using proprietary BioMark software. We built generalized mixed effects models (GLM) using R 4.1.2 (R Development Core team) and the R packages *glmmTMB*^44^ and *car*^45^ to examine relationships between predictor and response variables for all statistical comparisons, unless explicitly stated otherwise. Visual inspection of the relationship between variables and the resulting residuals from untransformed models guided the selection of any variable transformations. This ensured appropriate model fit and adherence to statistical assumptions. Most analyses employed a repeated measures design, examining data for each time unit per mouse or other relevant unit as appropriate. We included random effects of trial and animal identity to account for non-independence of observations. We occasionally needed to simplify the random effects structure to avoid singular fits or failures of model convergence. We report all means ± standard error (s.e.m.) unless explicitly stated otherwise. Results were considered statistically significant when *p* < 0.05. Full details of statistical analyses, including model formulae and fixed and random effect structures, are provided in the analysis code. Figures were created in R using the package *ggplot2*^46^, Inkscape 1.3.2, and TimeTree software^47^.

## RESULTS

We characterized circadian activity, space use, and social behavior profiles in replicate mixed-sex groups of *M. pahari* (PAH) mice in a large custom-built arena and compared them to C57BL/6J (B6) (n = 3 trials per genotype). Sixteen RFID antennae monitored the spatiotemporal activity patterns of mice (n = 4 males, n = 4 females per trial) over a five-day period (**Figs. 1C, S1A**). Our analyses focused on the comprehensive RFID dataset for the circadian, spatial, and social analyses (1,022,158 total RFID reads).

### Mus pahari exhibits more strongly stereotyped nocturnal circadian activity

Actograms revealed highly stereotyped circadian activity patterns in PAH mice, with transitions between antenna (hereafter ‘activity’) occurring almost exclusively during the lights off phase (1200-2200h) of the light-dark cycle (**Figs. 2A, S2, S3**). PAH mice exhibited a significantly higher percentage of dark cycle transition events compared to B6 mice (PAH: 98.5 ± 0.7%, B6: 76.6 ± 2%; **Fig. 2B**). We detected sex differences in B6 mice, but not in PAH mice, with B6 females showing a greater percentage of dark cycle transitions than males (**Fig. 2B**). Notably, B6 males showed a lower percentage of dark cycle activity due to increased light cycle transitions, indicating more flexible timing of activity throughout the 24-hour cycle (**Figs. 2B, S2**).

**Figure 2.**
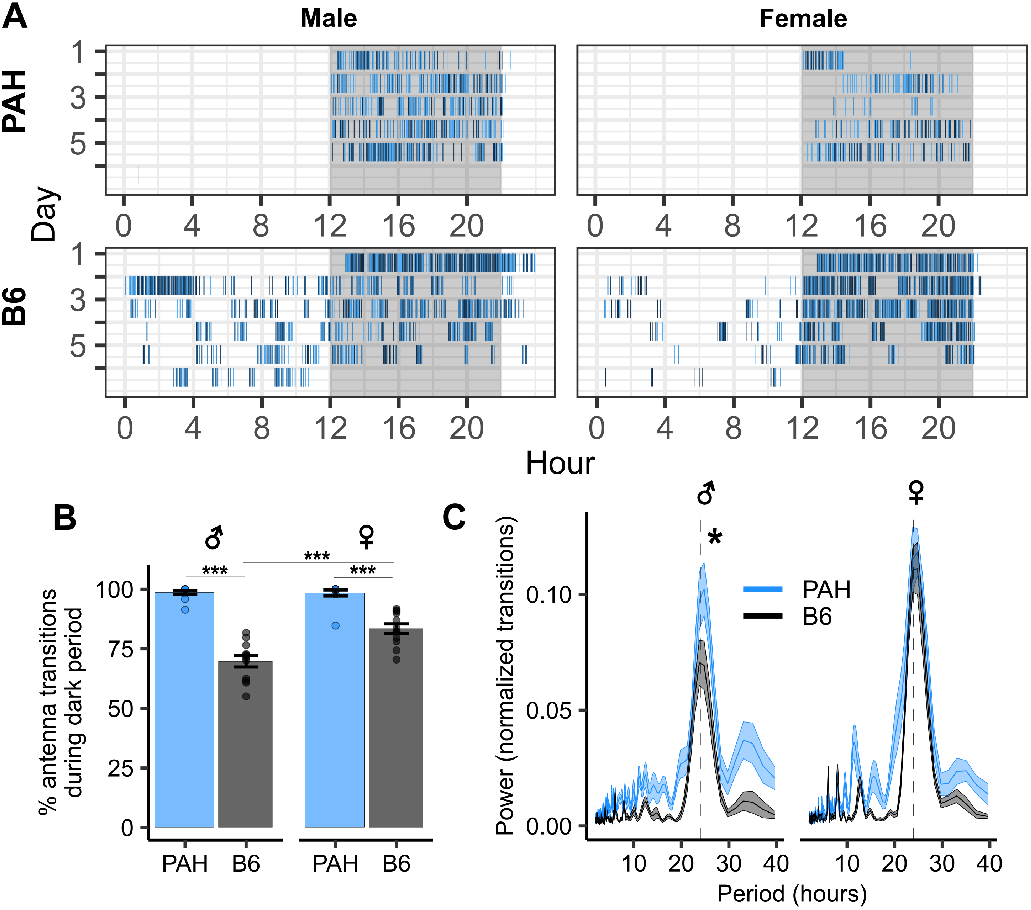
*Mus pahari* displays highly stereotyped dark-cycle circadian activity compared to *Mus musculus*. **A**. Representative actograms showing movement bouts from antenna transition events for individual male and female mice from each genotype. Colored ticks indicate transition events with color representing specific antenna locations. Shaded regions indicate the dark period (1200-2200h). Full trials for all individuals are presented in **Fig. S2. B**. Percentage of antennae transition events occurring during the dark period (1200-2200h) for males and females of both genotypes. **C**. Lomb-Scargle periodogram analysis of rhythmicity of movement bouts. Lines represent mean power spectra showing pronounced peaks around the 24-hour period (dotted line) for all groups. Higher peaks indicate stronger circadian rhythmicity. Despite similar period lengths (male peak period, PAH: 24.7 ± 0.17 hours, B6: 23.8 ± 0.3 hours; GLM_genotype_: *p* = 0.068), PAH males exhibited significantly stronger circadian rhythmicity compared to B6 males (male peak power, GLM_genotype_: *p* = 0.044). Female mice showed similar period lengths (female peak period, GLM_genotype_: *p* > 0.1, PAH: 24.4 ± 0.33, B6: 24.3 ± 0.19) and rhythm strength (female peak power, GLM_genotype_: *p* > 0.1) across species. B6 mice exhibited a significant sex difference in rhythm strength (B6 peak power, GLM_sex_: *p* = 0.006) not observed in PAH mice (PAH peak power, GLM_sex_: *p* > 0.1). Data are presented as mean ± s.e.m. *p* ≤ 0.05*, *p* ≤ 0.01**, *p* ≤ 0.001***. *P*-values extracted from generalized linear mixed effects models unless otherwise noted.

To further characterize circadian activity, we performed a Lomb-Scargle periodogram analysis on normalized antennae transition event data across all days of the experiment. This analysis allows for the identification of dominant periodicities in time series data even in the presence of missing or unevenly sampled data points, making it suitable for our antennae transition event data set. This analysis revealed strong 24-hour periodicity in both strains (PAH: 24.5 ± 0.2, B6: 24.1 ± 0.2), with a significant sex difference in B6 mice (males: 23.8 ± 0.3, females: 24.3 ± 0.2; **Fig. 2C**). Additionally, B6 males exhibited a significantly weaker circadian rhythm compared to PAH (*z* = −2.02, *p* = 0.044), as measured by the peak period power.

Our findings demonstrate that while both strains exhibit strong 24-hour circadian rhythms, PAH mice show more stereotyped activity confined to the dark phase compared to B6 mice, contrary to Marshall’s (1977) observations of recently wild-caught individuals. The strict nocturnal behavior of PAH mice suggests a more rigid adherence to light-dark cues, which may reflect adaptations to their natural ecological niche or reflect some aspect of their adaptation to the lab environment. Additionally, the sex difference observed in B6 mice, but not in PAH mice, suggests that *M. pahari* exhibits more canalized circadian behaviors that are less influenced by sex-specific factors in this strain. How this might relate to the evolutionary history of the two species versus the domestication of the inbred strains is unclear.

### Mus pahari exhibits reduced spatial exploration compared to Mus musculus

PAH exhibited significantly reduced exploration compared to B6 (**Figs. 3A, S3, S4**). While exploration of the arena was similar between B6 and PAH mice during the first hour of the trial, PAH mice exhibited slower exploration by the second hour (**Fig. 3B**). Strikingly, all B6 mice (24/24) explored all of the antenna monitored spaces over 5 days, whereas only 54.1% (13/24) of PAH mice were detected at every antenna at least once over the trial period (**Fig. S4B**). Notably, PAH mice exhibited sex differences in exploration not observed in B6. PAH females explored at a faster rate compared to PAH males, visiting significantly more antennae by the final day of the trial (females: 15.4 ± 0.3, males: 14.5 ± 0.4; **Fig. S4B**).

**Figure 3.**
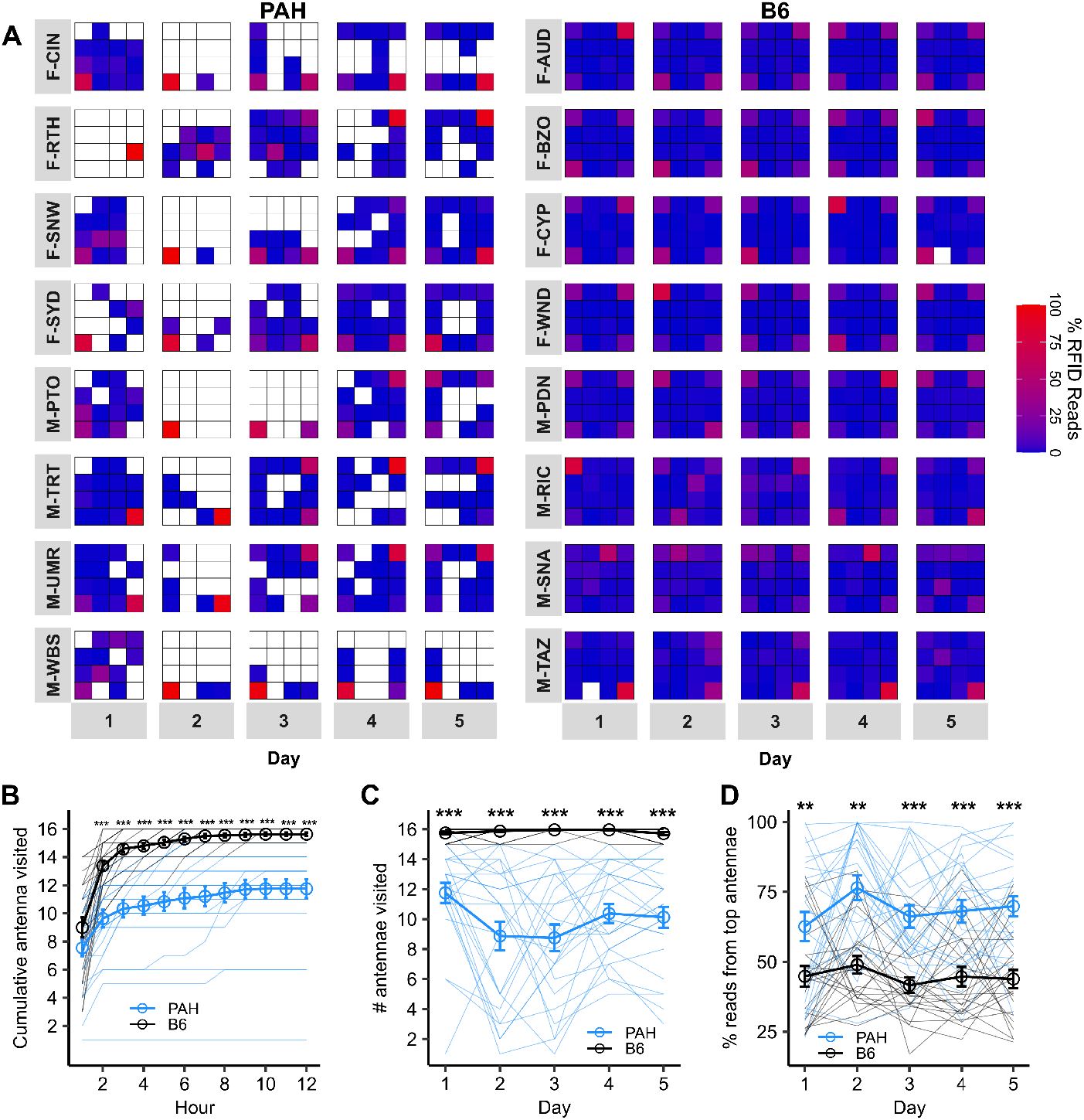
*Mus pahari* exhibit restricted exploration and enhanced spatial fidelity compared to *Mus musculus*. **A**. Semi-natural arena spatial occupancy heatmaps for *M. pahari* (PAH) and C57BL/6J (B6) across 5-day behavior trials (one trial per genotype shown). Individual male and female mice are indicated by M and F followed by a unique 3-letter identification code (e.g. F-CIN). The spatial layout of the 16 heatmap tiles corresponds to the layout of the 16 RFID antennae shown in **Fig. 1C**. Heatmap colors represent the percentage of an animal’s daily RFID reads at a particular antennae location, with warmer colors indicating higher occupancy (see **Fig. S4** for remaining trial data). **B**. Cumulative number of antennae explored during the first 12 hours of the trial. **C**. Number of unique RFID antennae visited per day. **D**. Daily spatial fidelity quantified by the percentage of daily RFID reads at an individual’s most frequented antenna per day. Data are presented as mean ± s.e.m. n = 24 per genotype. Solid lines represent individual subject data. *p* ≤ 0.05*, *p* ≤ 0.01**, *p* ≤ 0.001***. *P*-values extracted from generalized linear mixed effects models unless otherwise noted.

PAH mice visited significantly fewer antennae per day (males: 9.4 ± 0.5, females: 10.5 ± 0.5) compared to B6, which consistently visited nearly all antennae daily (averages) (**Fig. 3C**). In fact, there was only a single day that one PAH mouse visited all 16 antennas out of 120 total individual observation days across all trials and mice (i.e., 8 total mice over 5 days = 40 days per trials giving 120 days across all three replicates). In contrast, B6 visited all antenna on the vast majority of days (103/120 total individual observation days) and the fewest antennas visited by a B6 in a single day was 15 (17 out of 120 days). This restricted exploration pattern in PAH mice suggests a fundamentally different approach to spatial utilization compared to B6 mice. In addition to lower overall exploration, PAH of both sexes showed increased spatial fidelity to specific arena locations they did visit, quantified as the percentage of an individual’s total daily RFID reads derived from their most frequented antenna on that day, compared to B6 mice (males: *p* = 0.001, females: *p* = 0.003; **Fig. 3D**).

Movement patterns were also reflected in differential visitation of the four resource zones present in each trial. PAH initially visited fewer resource zones but gradually increased visitation over the trial. By days 4-5, PAH males, but not females, approached B6 levels of resource zone visitation (**Fig. S4C**). PAH mice showed strong initial resource zone bias towards their most frequented zone per day (>80% of zone reads on days 1-2), which decreased over time, suggesting a shift from exclusive to more distributed zone use (**Fig. S4D**). In contrast, B6 mice displayed relatively lower bias toward their most frequented zone throughout the trial (males: 57 ± 2.4%, females: 47.2 ± 1.7%; note that 25% would be random zone visitation), indicating a more balanced utilization of all available resource zones from the start (**Fig. S4D**). We next examined the degree to which zones were occupied by a specific male or female occupant compared to other individuals of the same sex by quantifying the percentage of the total male or female reads within a zone derived from the zone’s top occupant (relative to other same-sex competitors) per day (**Fig. S4E**). Consistent with initial low movement patterns, zones in PAH trials initially exhibited nearly exclusive occupation by a single male that decreased over time, while B6 male occupation patterns was lesser and stable across days. Zones in B6 trials were less exclusive to a single female compared to PAH trials.

### Enhanced social tolerance and male-male gregariousness in Mus pahari

We quantified social behavior by estimating the time mice spent overlapping in time at the RFID antennae using previously established methods^38^ (mean total hours of mouse observation time per trial ± s.e.m., PAH: 257.8 ± 34.3; B6: 351.1 ± 15.6; **Fig. S5A**). Analysis of co-occurrence at RFID antennae allowed us to infer the onset and offset of group formation events and time spent in various social grouping configurations (**Fig. 4A**). Overall, the majority of social grouping time occurred at antennas within the resource zones for both genotypes (PAH: 86.7%; B6: 73.4%).

**Figure 4.**
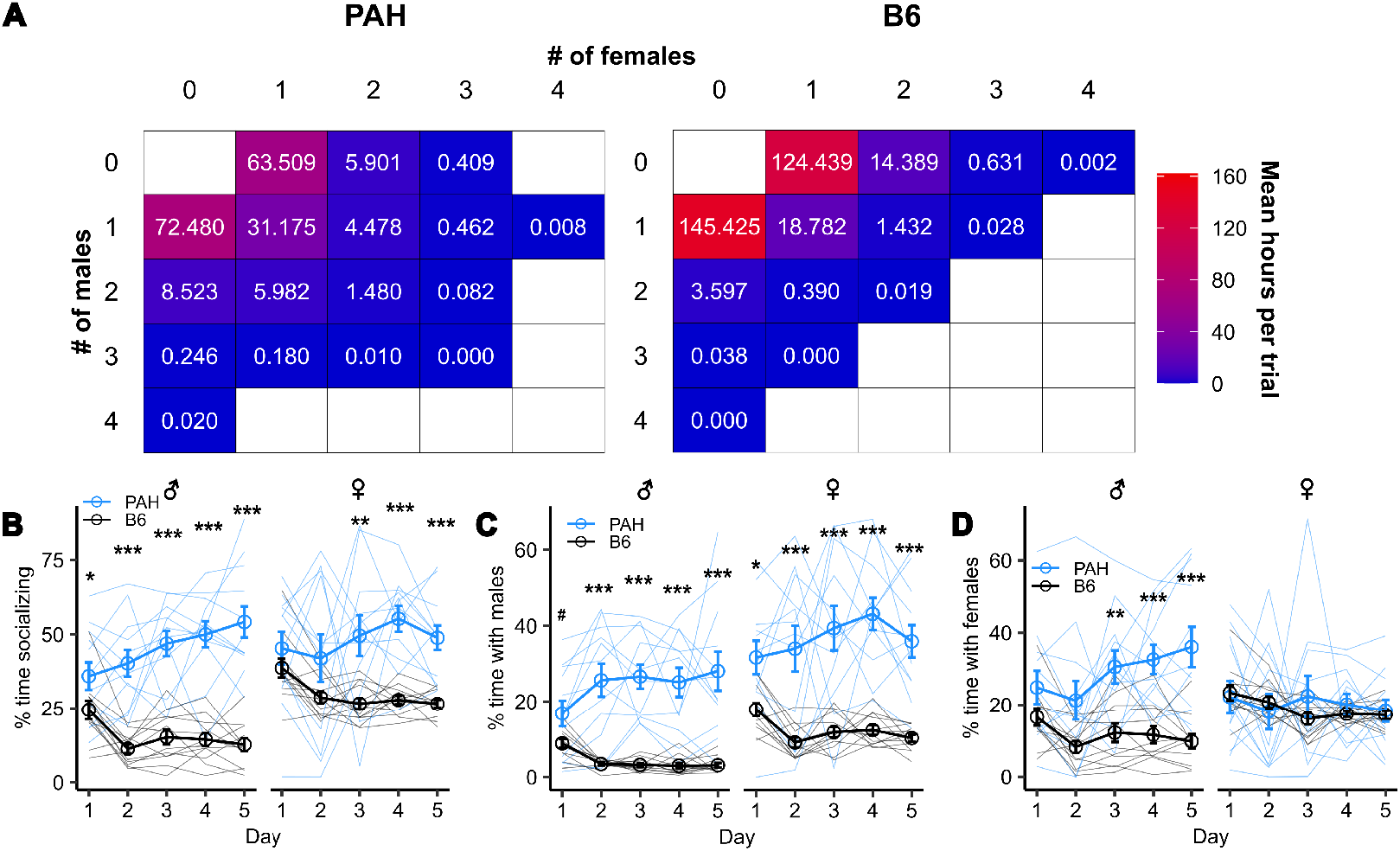
Enhanced social tolerance and male-male gregariousness in *Mus pahari* compared to *Mus musculus*. **A**. Heatmaps depicting the average time mice spent in various group compositions (combinations of males and females) per trial. Values indicate mean observed hours per trial. Warmer colors indicate more time spent in that group composition. **B**. Percentage of daily observed time spent socializing with at least one other conspecific across the 5-day trial period. **C**. Percentage of daily observed time spent in spatiotemporal association with males (unconditional of the presence of females). **D**. Percentage of daily observed time in spatiotemporal association with females (unconditional of the presence of males). Data are presented as mean ± s.e.m. Solid lines represent individual subject data. *p* ≤ 0.1^#^, *p* ≤ 0.05*, *p* ≤ 0.01**, *p* ≤ 0.001***. *P*-values extracted from generalized linear mixed effects models unless otherwise noted.

While both species spent the majority of their overall observed time alone, consistent with prior reports of B6 living in outdoor enclosures^38^, PAH demonstrated significantly higher sociality (mean daily percent time socializing ± s.e.m., PAH: 46.9 ± 1.7%; B6: 22.7 ± 1.0%; **Fig. 4B**). Notably, PAH mice increased the time spent in a group over the trial duration, contrasting with increasing time alone in B6 (*z* = −4.82, *p* < 0.001). By Day 5, PAH males spent 54.2 ± 5.2% of their observed time in social groups, compared to 12.7 ± 2.2% for B6 males. Intriguingly, while B6 females showed higher socialization than males (B6: *z* = 7.1, *p* < 0.001), PAH exhibited no significant sex difference in sociality (*p* = 0.615; **Fig. 4B**). These findings suggest a strong bias towards social interaction in PAH mice compared to B6 mice under semi-natural conditions.

We next considered how same- and opposite-sex social interactions differed between the species. PAH males exhibited a striking increase in overall time spent with other males over the course of the trial, while B6 males maintained relatively low-levels of male-male association (**Figs. 4C**). Additionally, PAH mice had longer multi-male spatiotemporal association bouts, suggesting the males tended to engage in more prolonged social interactions (**Fig. S5B**). PAH males also showed a gradual increase in time spent with females across days, contrasting with the more stable patterns observed in B6 (**Figs. 4D**). Female-female associations remained consistent across both strains (**Figs. 4D, S5C**).

## DISCUSSION

Our study provides novel insights into the spatial, circadian and social dynamics of an inbred wild-derived strain of *Mus pahari* (PAH) in comparison to a common lab strain of *Mus musculus domesticus* (B6). The results demonstrate that PAH mice exhibit distinct behavioral patterns, including highly stereotyped circadian activity, reduced spatial exploration, and enhanced male-male social tolerance in a small semi-natural indoor enclosure. These findings contribute to our understanding of phenotypic diversity within the *Mus* genus, highlighting social behavioral diversity among species.

Given the dominance of house mice in biomedicine, there are surprisingly few studies characterizing and comparing the behavioral diversity present across the *Mus* genus, either in the wild or in captive inbred strains and species. There are numerous reasons for this, including the technical challenges associated with studying wild populations of small rodents in their natural habitats, the geographic distribution of ancestral house mouse populations in the Eurasian steppe and middle east, and the effort of obtaining wild samples from species of *Mus* across the globe and adapting them to successfully breed in laboratory colonies. The existence of an inbred strain of *Mus* pahari provides a valuable opportunity to investigate the evolutionary history of behavioral phenotypes within *Mus*. Work that compares strains within and between species of *Mus* in similar common-garden assays has the potential to reveal how diverse behaviors have evolved across the genus. The work on non-*Mus musculus* species to date has focused on *Mus spretus* and *Mus spicilegus*. Future work that compares more species simultaneously has the potential to provide new insights into the evolution and biology of mice. Many understudied species of *Mus* are found in Africa and Asia offering opportunities for researchers in those locations to potentially work with those species in the lab. Comparisons of behavior to widely studied lab mouse strains, such as C57BL/6J offer an opportunity to use a standardized reference point for assays even if logistical constraints prevent individual labs from studying multiple non-*Mus musculus* species simultaneously.

The circadian movement patterns of PAH mice were highly stereotyped and tightly coupled to the dark cycle, contrasting with the more flexible movement patterns observed in B6 mice. While B6 frequently moved between antennas during the light phase of the light-dark cycle, there was almost no movement around the enclosure by PAH mice when the lights were on. It is important to emphasize that while movement between antennas unequivocally demonstrates activity, a lack of movement between antennae does not necessarily indicate that mice are not moving or otherwise inactive. For example, during the light phase mice could remain within a resource zone and sleep or they could remain and eat, socialize, or use a running wheel. Indeed, our observations of the mice suggested that PAH individuals were frequently engaging in some level of activity during the light phase of the light-dark cycle consistent with earlier anecdotal observations made by Marshall of wild *M. pahari*^24^. Nevertheless, the stark differences between genotypes in circadian rhythms in terms of movement around the enclosures highlight the potential for species-specific adaptations in circadian biology within the *Mus* genus.

Social interactions and space use are intimately linked and we observed differences in both between PAH and B6. PAH mice visited significantly fewer antennae and resource zones per day and showed higher spatial fidelity to specific areas. This could be interpreted as PAH individuals showing either less exploratory behavior or social behaviors leading individuals to divide space into territories. We did not measure movement and exploration of solitary mice, so it is difficult to disentangle social interactions and patterns of space use. However, the social behavior data suggest that increased territoriality is not present in PAH compared to B6 mice.

In contrast to the more solitary nature of male lab *M. m. domesticus* under semi-natural conditions (ref^38^ and **Fig. 4**), PAH males showed a marked increase in time spent with other males over the course of the trial, refuting a territorial hypothesis for the patterns of reduced spatial exploration. This was further evidenced by longer multi-male social association events in PAH trials compared to B6 trials. These patterns of increased male-male tolerance and group living in *M. pahari* are particularly interesting from an evolutionary perspective. While we cannot speculate on the ecological factors driving these behaviors without data from wild populations of *M. pahari*, the stark social differences from inbred *M. m. domesticus* reported here and from published studies on other *Mus* species^18^ suggests a strong variation in social strategies and organization within the *Mus* genus. Similar increased male social tolerance has been observed in other group living rodent species^48^, such as naked mole-rats^49^ and Cape Ground Squirrels^50^, although such tolerance is species specific and can vary with kinship status, and neither species directly matches the social organization reported here for *M. pahari*. The behavioral variation apparent within *Mus* provides an excellent opportunity for comparative genetic and neurobiological investigations into the mechanisms underlying diverse social strategies.

## Acknowledgements

We thank Dr. Matthew N. Zipple for helpful feedback on the manuscript. We thank Summer Hardy, Kwynn Guess, and Daniel Kuo for their assistance in turning over the mesocosm arena between trials.

## Ethics Approval

All experimental procedures were approved by the Cornell University Institutional Animal Care and Use Committee (IACUC: Protocol #2015-0060).

## Data and code availability

All tabulated data files and code will be made publicly available on GitHub upon publication of the final peer-reviewed article (https://github.com/calebvogt/2021_8x8_PAHvsC57). During the preprint stage, data and analysis scripts are available from the corresponding author upon reasonable request.

## SUPPLEMENTARY FIGURES

**Figure S1.**
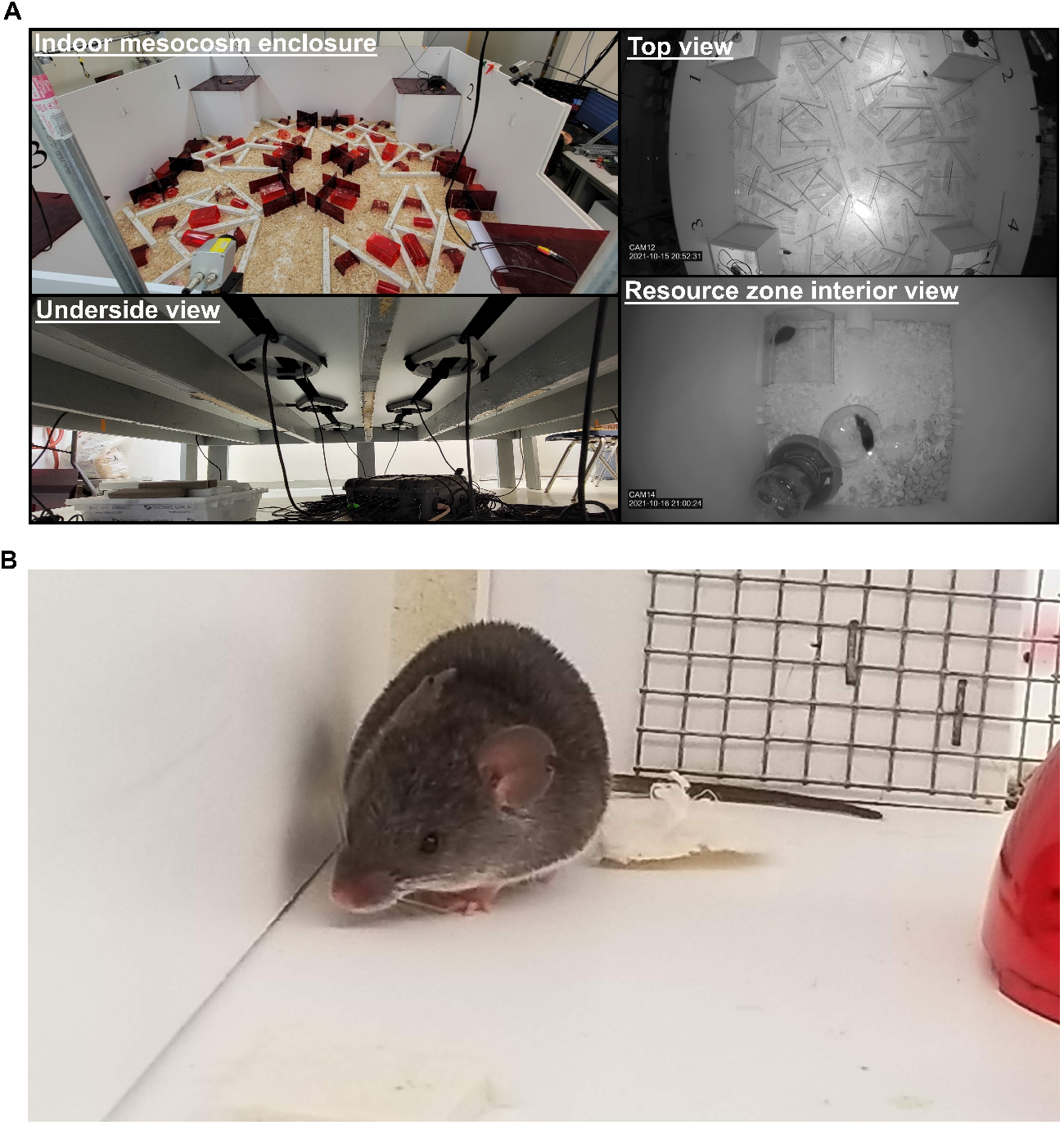
Indoor mesocosm enclosure setup. **A**. Images of the experimental setup: (Top left) Close-up view of the arena floor and walls, illustrating the experimental environment during the lights on period. (Top Right) Overhead view of the complete indoor mesocosm enclosure showing the 2.4 × 2.4 m arena with four resource zones in the corners. (Bottom left) Arrangement of the 16 RFID antennae mounted beneath the floor of the arena, demonstrating the comprehensive spatial coverage for tracking mouse movements. (Bottom right) Interior view of a resource zone containing enrichment materials, running wheel, and *ad libitum* food and water available to mice during the 5-day trials. **B**. Representative image of a *M. pahari* (PAH/EiJ) mouse, showing the distinctive shrew-like morphological features including elongated snout, large ears, and blueish-gray velvet spiny coat that differentiate this species from *M. musculus domesticus*.

**Figure S2.**
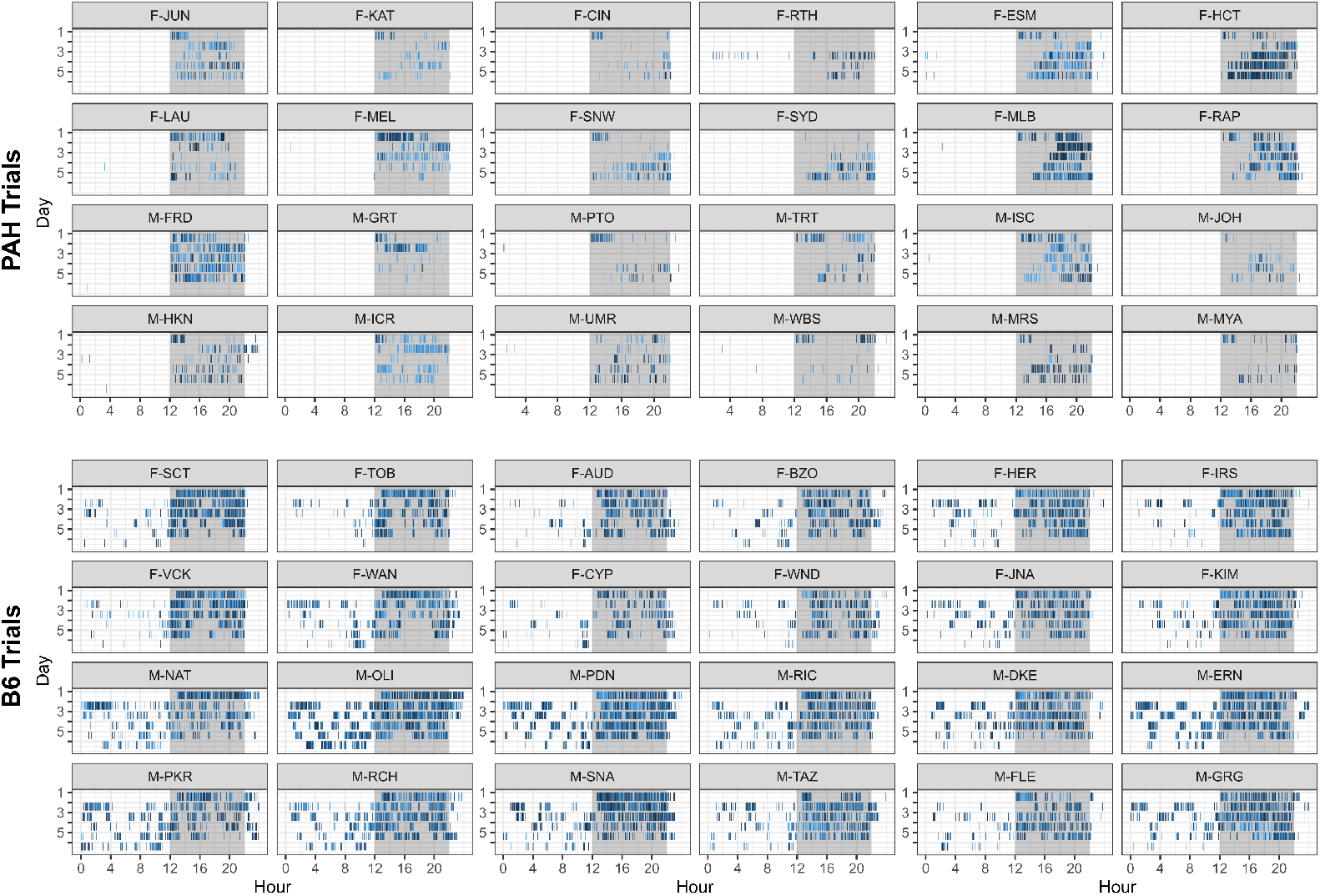
Circadian actograms for PAH and B6 trials. Full trial circadian actogram data across days for PAH and B6 mice. Shaded regions represent the dark period (1200-2200h). Colored tick marks indicate antenna transition events, with colors denoting specific antenna identity.

**Figure S3.**
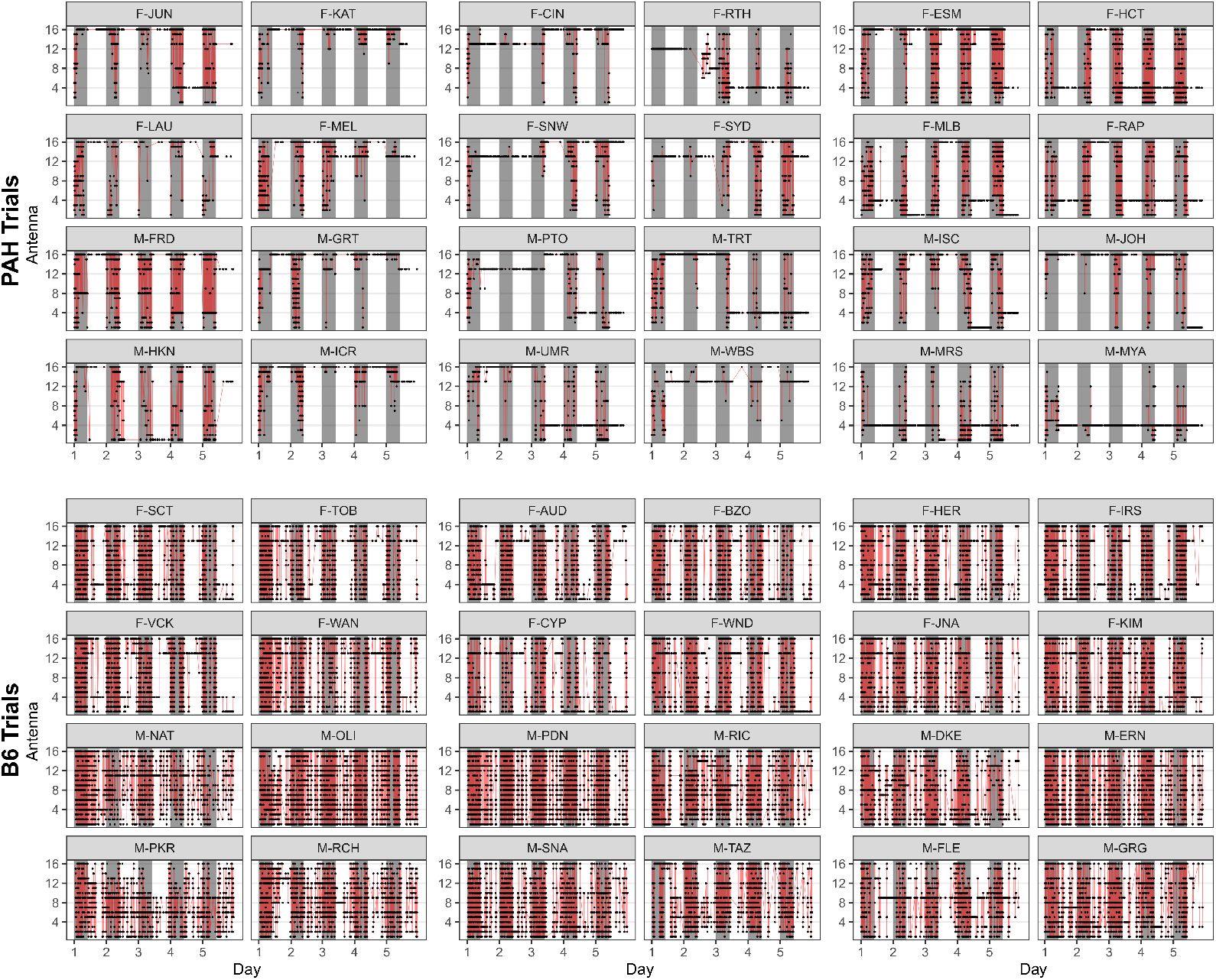
Spatiotemporal patterns of RFID reads across genotypes reveal distinct daily activity profiles. Full trial data of individual RFID read events at the 16 RFID antennae in the arena for PAH and B6 mice (n = 3 trials per genotype). Black dots indicate individual RFID reads, while red lines trace transitions between sequential antenna reads, illustrating patterns of activity and movement. Sex and individual are designated by 4-letter codes (e.g. “F-JUN”). Shaded gray regions indicate the duration of the dark cycle between 1200-2200h.

**Figure S4.**
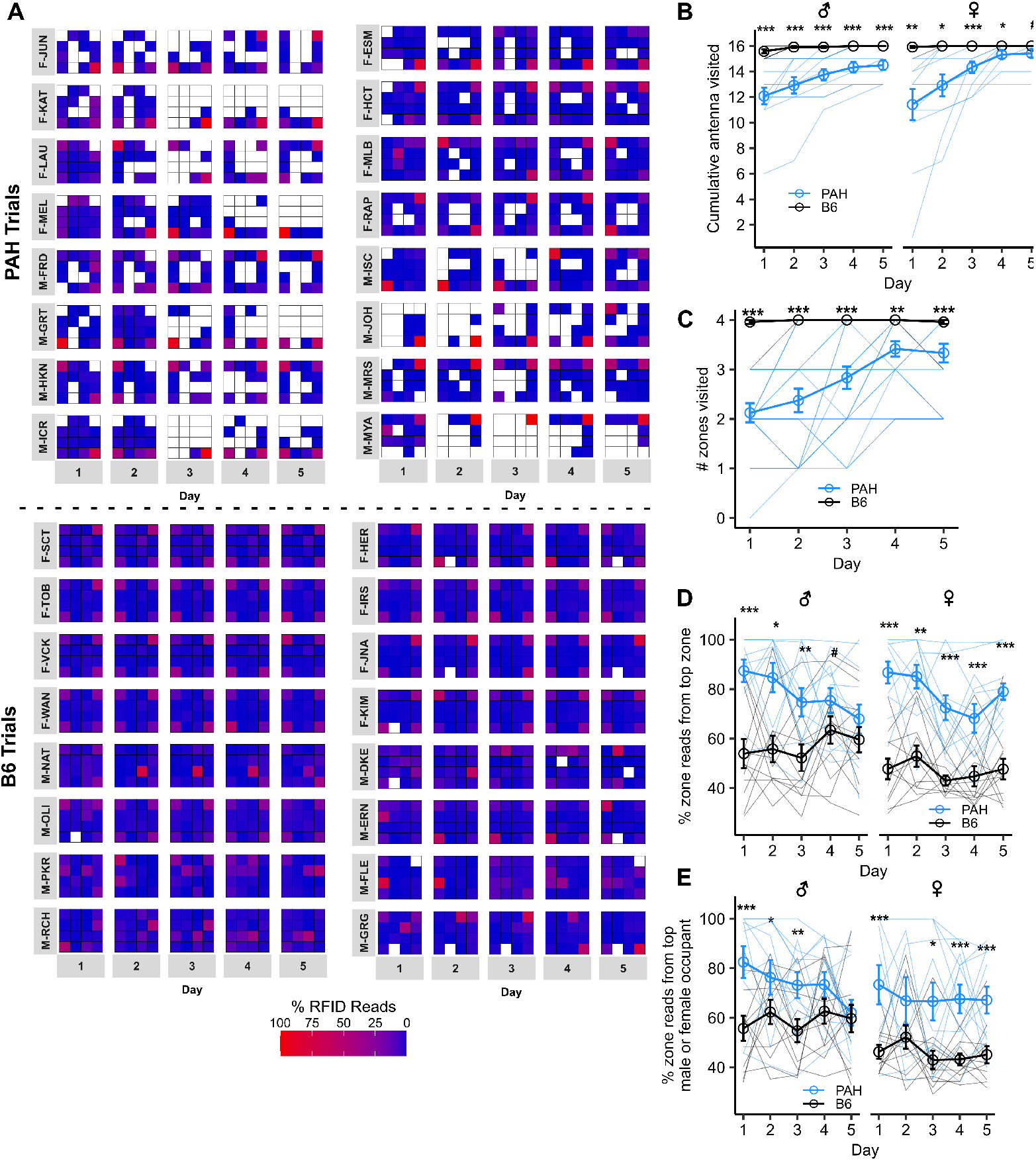
Spatial exploration and resource utilization patterns in *Mus pahari* and *Mus musculus*. **A**. Full trial spatial occupancy heatmaps across 5-day trials (two trials per genotype shown; see **Fig. 2A** for remaining trial). Individual male and female mice are indicated by sex (M/F) and a unique 3-letter identification code. Colors represent the percentage of an animal’s daily RFID reads at a particular antenna location, with warmer colors indicating higher occupancy. **B**. Cumulative number of unique antennae visited over the 5-day trial period for males and females of both genotypes. **C**. Number of unique resource zones visited per day by individual mice. **D**. Resource zone fidelity measured by the percentage of zone RFID reads at an individual’s most frequented zone per day. **E**. Percentage of a resource zone’s total daily reads which were derived from either the top male or top female occupant relative to all other same-sex RFID reads in that zone (n = 4 zones in n = 3 trials per genotype). Solid lines represent unique zones in each trial. Data are presented as mean ± s.e.m. Solid lines represent individual subject data unless otherwise indicated. *p* ≤ 0.1^#^, *p* ≤ 0.05*, *p* ≤ 0.01**, *p* ≤ 0.001***. *P*-values extracted from generalized linear mixed effects models unless otherwise noted.

**Figure S5.**
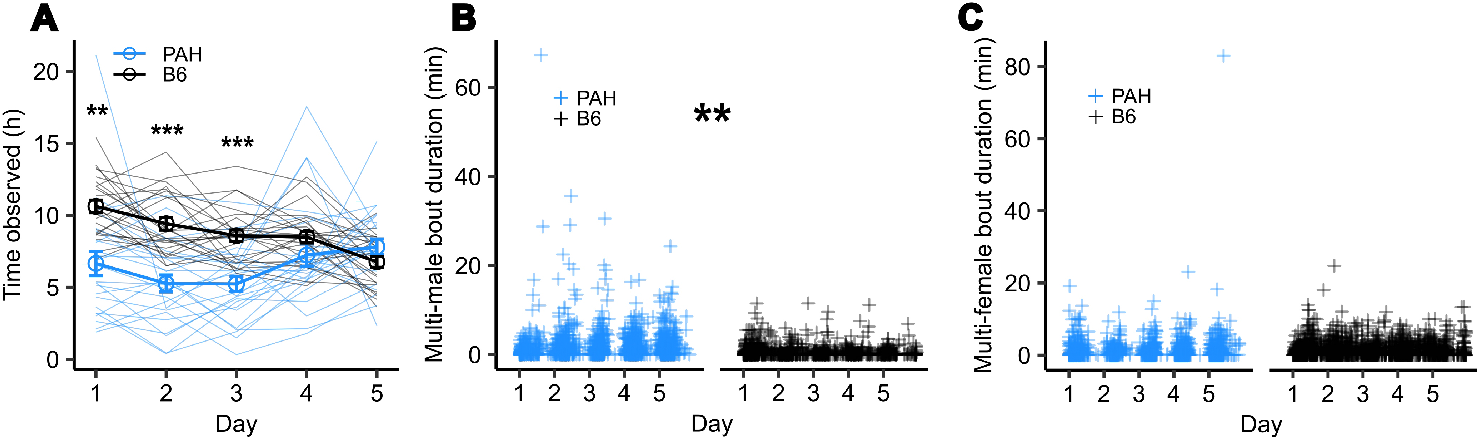
Enhanced male-male social contact in *Mus pahari* compared to *Mus musculus*. **A**. Total estimated time individuals were observed as occurring in spatiotemporal association with RFID antenna locations, either alone or in social groups, across the 5-day trial period. **B**. Duration of multi-male social spatiotemporal association events across trials (n = 1646 multi-female PAH bouts, n = 1318 multi-female B6 bouts). Plus signs represent individual association events. **C**. Duration of multi-female social spatiotemporal association events across trials (n = 1417 multi-female PAH bouts, n = 2461 multi-female B6 bouts). Plus signs represent individual association events. Data are presented as mean ± s.e.m. Solid lines represent individual subject data unless otherwise indicated. *p* ≤ 0.05*, *p* ≤ 0.01**, *p* ≤ 0.001***. *P*-values extracted from generalized linear mixed effects models unless otherwise noted.

